# Oxygenated hemoglobin signal provides greater predictive performance of experimental condition than de-oxygenated

**DOI:** 10.1101/2021.11.19.469225

**Authors:** Robert Luke, Maureen J Shader, Alexandre Gramfort, Eric Larson, Adrian KC Lee, David McAlpine

## Abstract

Continuous-wave functional near-infrared spectroscopy (fNIRS) neuroimaging provides an estimate of relative changes in oxygenated and de-oxygenated hemoglobin content, from which regional neural activity is inferred. The relation between both signals is governed by neurovascular coupling mechanisms. However, the magnitude of concentration changes and the contribution of noise sources to each chromophore is unique. Subsequently, it is not apparent if either chromophore signal practically provides greater information about the underlying neural state and relation to an experimental condition. To assess this question objectively, we applied a machine-learning approach to four datasets and evaluated which hemoglobin signal best differentiated between experimental conditions. To further ensure the objective nature of the analysis, the algorithm utilized all samples from the epoched data rather than pre-selected features. Regardless of experimental task, brain region, or stimulus, the oxygenated hemoglobin signal was better able to differentiate between conditions than the de-oxygenated signal. Incorporating both signals into the analysis provided no additional improvement over oxygenated hemoglobin alone. These results indicate that oxyhemoglobin is the most informative fNIRS signal in relation to experimental condition.

## Introduction

Functional near-infrared spectroscopy (fNIRS) is a non-invasive neuroimaging tool that uses near-infrared light to infer the relative change in hemoglobin concentration in cortical blood flow. Oxygenated hemoglobin (HbO) and de-oxygenated hemoglobin (HbR) have distinct absorption spectra within the near-infrared range, making it possible to establish concentration changes for HbO and HbR chromophores separately, as well as total hemoglobin—the sum of the oxygenated and de-oxygenated components (HbT) (Ferrari et al., 2004). fNIRS provides a number of advantages over other neuroimaging techniques. Besides its ability to differentiate relative changes in the concentration of oxy- and de-oxygenated hemoglobin, fNIRS has a higher sampling rate and is less sensitive to motion artifacts compared to fMRI. However, scattering of the near-infrared light limits the penetration depth of the signal into cortical tissue and decreases the signal-to-noise for the neural signal of interest (Scholkmann and Wolf, 2013). Further, physiological noise arising from non-neural processes (e.g., Mayer waves, respiration, heart rate) can also contaminate the signal of interest (Kirilina et al., 2013; Luke et al., 2021b). Robust analysis techniques are therefore required to overcome the inherent limitations of fNIRS measurements. In particular, it has been acknowledged that there is no consensus as to which chromophore, or what combination or contrast of chromophores, is best suited for differentiating between different neural states across various fNIRS neuroscience applications (Herold et al., 2018; Kohl et al., 2020; Lloyd-Fox et al., 2010).

Each chromophore is differentially impacted by the multiple sources of noise in fNIRS signals. The oxygenated signal exhibits a greater amplitude than the de-oxygenated signal (Sato et al., 2016; Stangl et al., 2013) and has greater test-retest reliability (Plichta et al., 2006). Conversely, the de-oxygenated signal is less affected by extracranial physiological noise (Dravida et al., 2017; Heinzel et al., 2013; Kirilina et al., 2012; Obrig et al., 2000) and provides greater spatial selectivity (Cannestra et al., 2003; Plichta et al., 2007). The cortical contribution within total hemoglobin signals, however, may be significantly higher than both oxy- and de-oxygenated signals because total hemoglobin is far less sensitive to extracerebral contamination (Gagnon et al., 2012). A recent report discussing best practices in fNIRS studies recommends that researchers present a visualization of both chromophores—HbO and HbR—and the statistical outcomes associated with both and a justification if one of the two is not reported (Yücel et al., 2021). Presenting both chromophores provides information as to the temporal characteristics of hemoglobin changes, as well as the quality of the data (Tachtsidis and Scholkmann, 2016). While there are advantages to reporting both chromophores, it is unclear if the two chromophores provide independent information about neural activity, and if not, which chromophore provides the better signal from which to extract and infer the neural activity of interest.

Beyond fundamental neuroscience applications, fNIRS-based brain-computer interface (BCI) applications typically perform feature selection using oxy- and de-oxygenated chromophores and/or the linear combination of both (total hemoglobin). In general, either the mean or slope of the oxygenated signal is a more robust classifier than de-oxygenated or total hemoglobin for task-evoked neural activity (Gemignani and Gervain, 2021; Mihara et al., 2012; Naseer and Hong, 2013; Naseer et al., 2014). However, the manual selection of features based on the time course of the fNIRS signal may bias the results towards one chromophore, as the time courses of these signals differ in amplitude and latency (Scholkmann et al., 2014; Tam and Zouridakis, 2014). Further, the generalizability of these findings to different brain regions and tasks is unknown and this is of specific importance in fNIRS investigations as both systemic noise and evoked neural responses vary with task, location of optical sources and detectors on the scalp, and the type of stimulus presented (Yücel et al., 2017; Zhang et al., 2015). As such, a methodologically unbiased systematic investigation utilizing different experimental tasks, scalp locations, and stimuli is required to determine which chromophore(s) is/are best suited from which to infer the neural signature in fundamental neuroscience research and its applications.

Based on the intrinsic relationship between both chromophores, signal enhancement techniques have been developed to exploit this relationship to improve signal quality and reduce the impact of physiological noise (Cui et al., 2010). However, objective quantification of these methods, in terms of their performance to differentiate between experimental conditions based on brain signals (decoding), has not been evaluated. Since de-oxygenated signals suffer from poorer signal-to-noise ratios, it is possible that the inclusion of de-oxygenated hemoglobin would not improve classification performance above oxygenated hemoglobin alone, or that it could potentially introduce additional noise. Here, we assess decoding accuracy for each chromophore and combinations of chromophores, with and without anticorrelation enhancement methods applied, for fNIRS measurements using multiple sensory modalities of vision, hearing, and motor movement.

Our overall aim was to evaluate the informational content of each chromophore, and the combination of both chromophores, to task-evoked fNIRS measurements for multiple experimental paradigms, stimulus types, and brain locations. To this end, a machine-learning approach was used to quantify the ability of each signal component to differentiate between conditions. Machine-learning analysis utilizes a data-driven approach to determine the optimal combination of input features to classify the trials by condition. To provide an experimental-design agnostic evaluation of classification performance, the raw samples from epoched data were utilized in the decoder rather than manual selection of signal features, such as peak, slope, or mean value of the signal (Gemignani and Gervain, 2021; Naseer and Hong, 2015). By utilizing a stratified k-fold cross-validation scheme, the performance of each signal component to differentiate between neural responses to different experimental stimuli was evaluated.

## Methods

We quantified the performance of different signal chromophores and signal processing to differentiate between experimental conditions from four datasets, three of which are publicly available. To ensure the conclusions from this study were generalizable, the datasets were selected to cover a variety of brain regions (occipital, motor cortex, auditory cortex), different task types (passive vs. active responses), and different stimulus modalities (auditory and visual). Further, canonical experimental designs were included, such as finger tapping and visual checkerboards, as well as more challenging experimental tasks for the fNIRS modality, such as detecting auditory-evoked responses (which is challenging due to the neural structures of interest lying relatively distant from the location of optodes on the scalp).

### Datasets

Each dataset is briefly described here in turn. Complete details of the experimental setup and recording parameters are provided in the associated publications (Luke et al., 2021a; Luke and McAlpine, 2021; Shader et al., 2021). All data were collected using NIRx NIRScout devices, although the exact device, optode placement, and sample rate varied across each dataset. All measurements included an additional eight source-detector pairs with a separation distance of 8 mm (short-distance channels) designed specifically to measure extracerebral changes in oxygenated and de-oxygenated hemoglobin. All experiments utilized a block design with randomized inter-stimulus-interval durations.

### Finger Tapping

In this active experimental task, five participants were asked to tap their thumb to fingers on the same hand when they heard an auditory cue (Luke and McAlpine, 2021). Participants used either their right or left hand only during any experimental block, and the hand used was indicated by a sound. If the sound was presented in the left earphone, participants tapped with their left hand, and similarly, if in the right earphone, participants tapped with their right hand. Participants tapped their thumb to fingers for the 5 sec duration of the sound. Each condition, either right- or left-handed tapping, was presented 30 times in a random order with a random inter-stimulus interval of 15-30 seconds. Optodes were placed over the motor cortex (Figure 1 top left). To evaluate model performance using this dataset, the ability to differentiate between left-hand and right-hand tapping based on the fNIRS measurement was evaluated.

**Figure 1:**
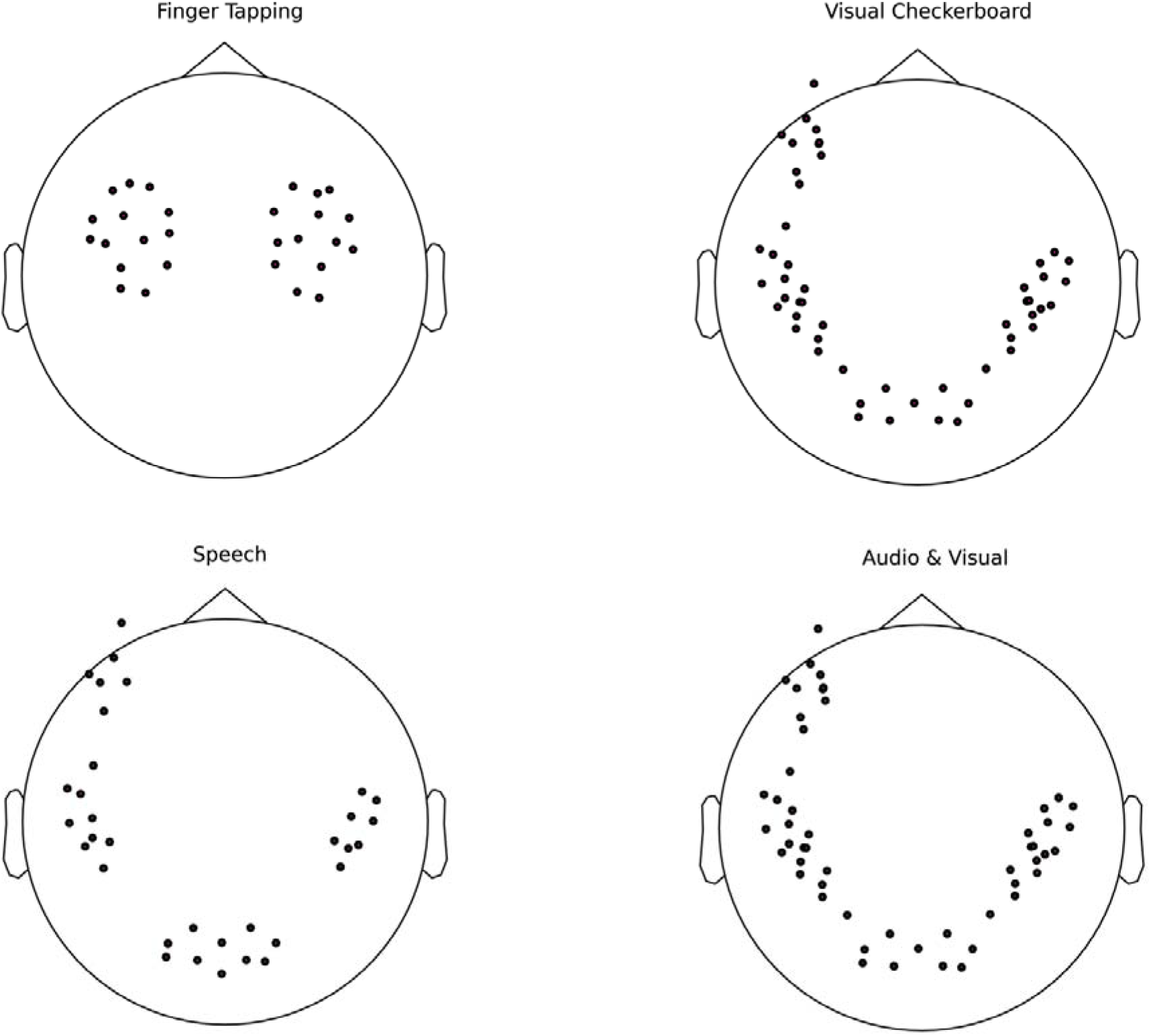
Channel placement for each dataset. Channel locations were defined as the midpoint between the source and detector optodes. The machine learning algorithm used data from all channels. Note that points outside the head circle indicate the channel was located below the vertical midpoint of the head.

### Visual Checkerboard

In this passive experimental task, six participants were asked to focus their vision on a computer monitor positioned one meter in front of them. The screen displayed either an oscillating black-and-white checkerboard pattern (condition) or a static grey cross (control) in the center of a black screen. The checkerboard oscillated at a rate of 4 Hz and each square had an edge length of approximately 2 cm. The stimulus duration was 5 sec with an inter-stimulus interval of 15-30 sec. Between stimulus presentations, the screen displayed only a grey cross, indiscernible from the cross (control) condition, thus no difference in neural activity was expected for the cross condition. The checkerboard condition was repeated 20 times and the control condition ten times, with the order of conditions randomized. Optodes were placed over the left inferior frontal gyrus, visual cortex, and auditory cortices (Figure 1 top right). To evaluate model performance using this dataset, the ability to differentiate between the checkerboard and control condition based on the fNIRS measurement was evaluated.

### Auditory Speech

In this passive experimental task, seventeen participants were asked to watch a silent subtitled movie of their choice. Simultaneously, over headphones, they were presented with either speech, low-frequency noise, or silence in a randomized experimental order (Luke et al., 2021a). Each stimulus was presented for 5 sec with an inter-stimulus interval of 15-30 seconds, and each was repeated 20 times in a randomized order. Optodes were placed over the left inferior frontal gyrus, visual cortex, and auditory cortices (Figure 1 bottom left). To evaluate model performance using this dataset, the ability to differentiate between auditory speech and silence condition based on the fNIRS measurement was evaluated.

### Auditory vs Visual Speech

In this active experimental task, eight participants were asked to pay attention to a running story that was presented in short segments in a block paradigm. Each experimental block presented the story segment either as auditory only or visual only (a face presented on a monitor speaking the sentence, without sound) (Shader et al., 2021). The order of conditions was randomized per participant and 18 trials were presented of each condition; the average trial duration was 12.5 seconds. Optodes were placed over the left inferior frontal gyrus, visual cortex, and auditory cortices (Figure 1 bottom right). To evaluate model performance using this dataset, the ability to differentiate between the auditory-only and visual-only conditions based on the fNIRS measurement was evaluated.

### Analysis

All analyses were performed using MNE v0.24 (Gramfort et al., 2013; Gramfort et al., 2014), MNE-NIRS v0.1.1 (Luke et al., 2021a), and scikit-learn v1.0 (Pedregosa et al., 2011). The same pre-processing was applied to each dataset and a minimal processing pipeline was applied to each measurement. Several additional pre-processing steps were also evaluated. An example of the pipeline is provided on the MNE-NIRS web page.

After data collection, all data was converted to BIDS format (Gorgolewski et al., 2016) using MNE-BIDS (Appelhoff et al., 2019). The signal was first downsampled to 1.5 Hz, then converted to optical density, and then hemoglobin concentration using the modified Beer-Lambert Law with a differential pathlength of 6. A low-pass filter was applied to the data with a cut-off frequency of 0.6 Hz and transition bandwidth of 0.1 Hz. This signal was restructured into epochs from 5 seconds before to 30 seconds after stimulus onset. The stimulus duration was different across datasets, however, a fixed post-onset epoching time was selected to ensure analysis was consistent across all data. Each epoch was baseline corrected based on the pre-stimulus time segment and a linear detrend was applied to each epoch. Epochs with a peak-to-peak amplitude greater than 100 μM were rejected. The epoched data were utilized in the machine-learning procedure.

Two machine-learning metrics were extracted from each dataset. For qualitative analysis, a time-by-time decoding approach was used to view the decoding performance at each sample in the epoched data. Second, for quantitative analysis, a single spatio-temporal metric approach was used that simultaneously used all channels and time points to estimate the experimental condition. (1) A machine-learning procedure employing this time-by-time decoding approach consisted of first standardizing each feature, which amounts to scaling each data point for each channel by the mean and standard deviation from all epochs. After which, a logistic regression classifier was applied using the *liblinear* solver (Fan et al., 2008). (2) For the spatio-temporal approach, the data were scaled for each channel by the mean and standard deviation from all time points and epochs, after which they were vectorized to comply with the scikit-learn data structure, and a logistic regression classifier was applied using the *liblinear* solver. These approaches classify the data within, rather than across, subjects; the intra-subject analysis approach was taken due to factors such as hair and skin color that grossly affect inter-subject variability. For both approaches, a 5-fold cross-validation score was determined, providing a robust estimate of the model performance to unseen data. The area under receiver operating characteristic curve (ROC AUC) was used as the evaluation metric to generate a single value that encompasses both the true and false positive performance of the model. The average of the ROC AUC metric across cross-validation folds is reported.

To evaluate the effective performance of each signal component for differentiating between experimental conditions, various signal components were used as input to the machine learning algorithm, including 1) oxyhemoglobin (HbO) only, 2) de-oxyhemoglobin (HbR) only, 3) both oxy- and de-oxyhemoglobin (HbO & HbR), 4) both oxy- and de-oxyhemoglobin with negative correlation enhancement applied directly before the epoching procedure (Cui et al., 2010) (HbO & HbR w/ enhancement), and 5) total hemoglobin (HbT; I.e., HbO + HbR). Thus, for each component of the fNIRS signal, an estimate of the ability to differentiate between conditions was obtained. To compare the performance of HbO and HbR, a two-sided paired t-test was performed comparing the average performance per dataset.

## Results & Discussion

To determine which component of the fNIRS signal provided the greatest information of the brain state related to experimental condition, a machine-learning analysis was applied to various chromophore and combinations of chromophore signals. The ability to differentiate between conditions was above chance for all datasets and all fNIRS signal components. The decoding performance for the pre-stimulus period was at chance level for all datasets (Figure 2), demonstrating correct implementation of the decoding procedure.

**Figure 2:**
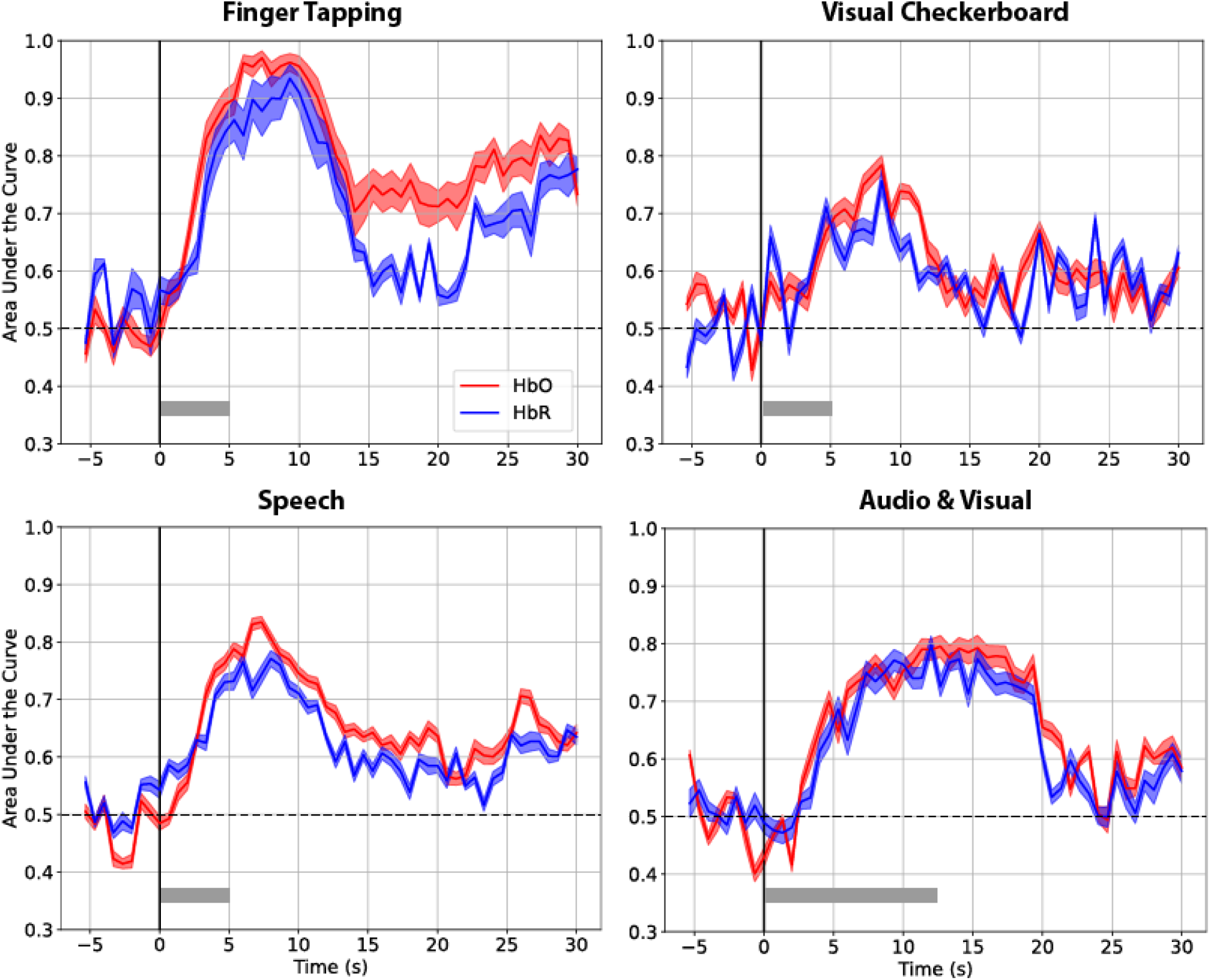
Time-by-time decoding performance for each dataset. Each curve represents the cross-validation receiver operating characteristic area under the curve scores averaged across subjects, and the error bars represent the standard error of the mean. The stimulus duration is indicated by the gray bar. The performance for different input signals to the machine learning algorithm are illustrated (color). Performance before stimulus onset (time = 0) is consistently at chance level (50%). Decoding performance improves after stimulus onset and peaks several seconds after stimulus offset.

The time-by-time decoding performance peaked within the first 10 seconds post stimulus-onset for finger tapping, checkerboard, and speech datasets, consistent with the 5-sec stimulus duration and known temporal characteristics of the hemodynamic response (Glover, 1999; Tak and Ye, 2014). Whereas the decoding performance for the auditory-visual dataset maintained its peak decoding performance for approximately 20 sec, consistent with the longer ~12.5 sec stimulus. The optimal decoding time was typically following the offset of the stimulus, and at or a few seconds after the peak in the oxyhemoglobin response (Figure 3). This illustrates that manual selection of signal features, such as signal value at the end of stimulus/task duration or windowed mean values over a stimulus duration, may not capture the optimal decoding performance, which peaks several seconds post stimulus offset.

**Figure 3:**
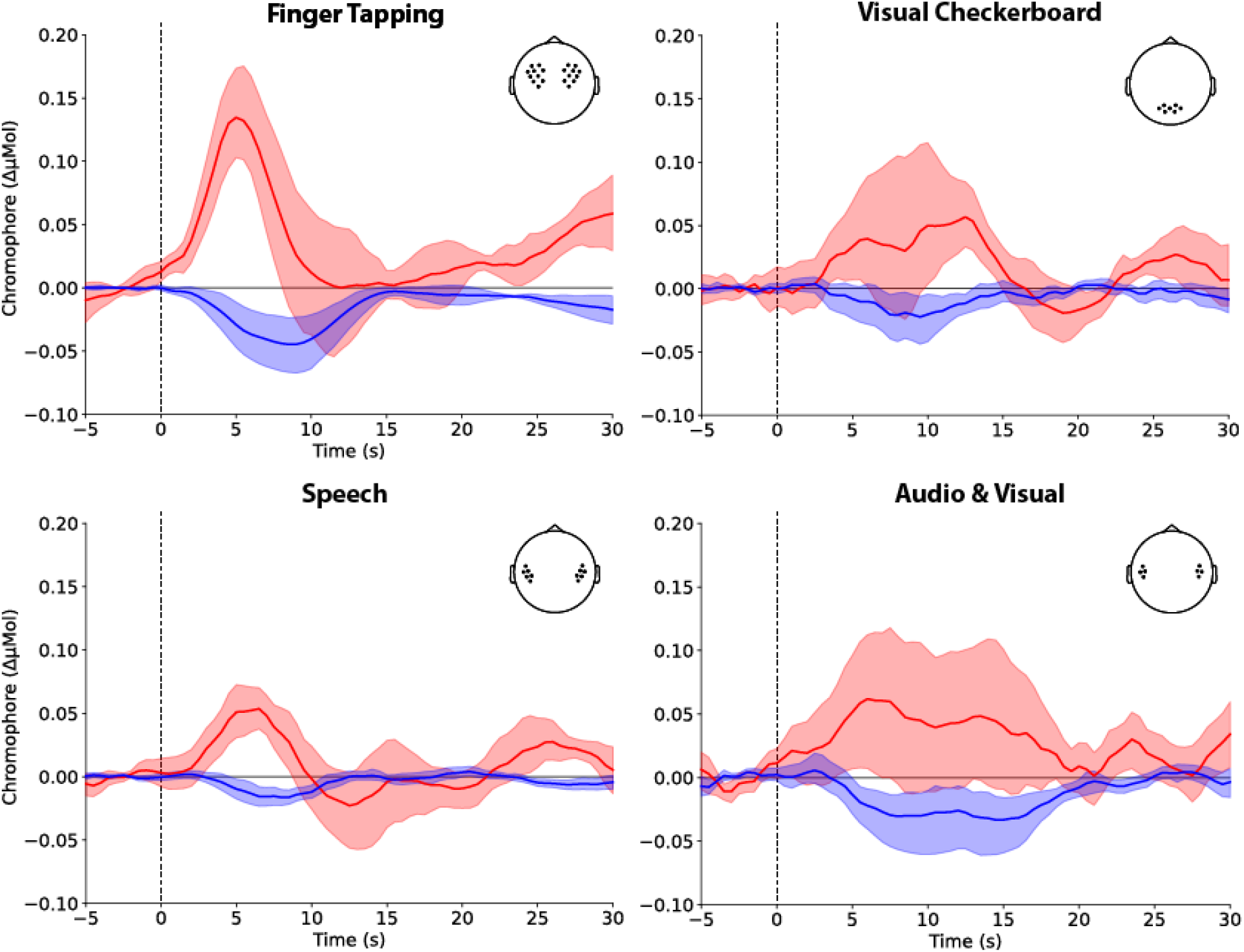
Grand average waveforms for a single condition and region of interest per dataset. Both oxygenated haemoglobin (red) and deoxygenated haemoglobin (blue) waveforms are shown. See the source manuscripts for each individual dataset for additional waveform figures. Note that for illustrative purposes the waveforms are taken from select regions of interest as illustrated in the inset. Whereas, the machine learning model used data from all available optode channels.

The decoding performance did not return to baseline within 30 seconds of the stimulus-onset for all datasets. This is likely due to the epoch length overlapping with the following stimulus presentation, with a 5-sec stimulus and minimum ISI of 15 sec; the next stimulus may occur as early as 20 seconds. However, the 30 second duration of post-stimulus epoching was selected to allow identical processing to be applied to each dataset.

The oxygenated hemoglobin signal provided the best decoding performance (Table 1), outperforming both the de-oxyhemoglobin and total hemoglobin signals. The mean ROC-AUC was greatest for the HbO data in all four datasets and was larger than the HbR data (M=7.7, SD=2.3), t(3)=5.6, p=.011. This result is consistent with BCI reports (Naseer and Hong, 2015), which suggest improved BCI performance for oxyhemoglobin signals, although it has not been directly addressed in a structured investigation such as this that investigates multiple stimulus types and brain regions.

**Table 1:** Decoding performance for fNIRS signals for the spatio-temporal analysis. Decoding performance was evaluated using the area under the curve of the receiver operating characteristic curve. The mean decoding performance (and standard deviation) is reported. A higher percentage indicates better performance. Performance was evaluated using oxygenated hemoglobin (HbO), de-oxygenated hemoglobin (HbR), total hemoglobin (HbT), both oxygenated hemoglobin and de-oxygenated hemoglobin (HbO & HbR), and both oxygenated hemoglobin and de-oxygenated hemoglobin with negative correlation enhancement (HbO & HbR w/ Enhancement). Results indicate that including de-oxygenated hemoglobin as input in addition to the oxygenated hemoglobin signal to the machine learning model provides no performance improvement in decoding between experimental conditions based on fNIRS neuroimaging signals.

|  | HbO | HbR | HbT | HbO & HbR | HbO & HbR w/<br>Enhancement |
| --- | --- | --- | --- | --- | --- |
| Tapping (N=5) | **89% (10)** | 85%<br>(15) | 82%<br>(15) | 88% (11) | **89% (13)** |
| Checkerboards (N=6) | **65% (20)** | 55%<br>(11) | 63%<br>(19) | 61% (19) | 64% (20) |
| Auditory Speech (N=17) | **72% (16)** | 61%<br>(20) | 68%<br>(12) | 67% (19) | 65% (21) |
| Auditory-Visual (N=8) | **76% (14)** | 70% | 74% | 74% (16) | 70% (18) |
|  |  | (20) | (16) |  |  |

Including both oxy- and de-oxygenated hemoglobin provided no additional decoding performance (Table 1). All analyzed signals, including the de-oxygenated signal alone, provided greater-than-chance decoding performance. Together, the data indicate that the information provided by de-oxygenated hemoglobin is not completely independent of that conveyed by the oxygenated hemoglobin signal, perhaps due to both signals reflecting the same underlying neurovascular coupling mechanism. When both oxy- and de-oxygenated hemoglobin signals were included as inputs, average performance of the model was slightly lower than when the HbO signal only was assessed. This reduction in performance is likely due to the increased dimensionality of the machine-learning problem. Similarly, a model applied to the total hemoglobin signal was no better than a model applied to the oxygenated signal alone, despite metrics based on total hemoglobin potentially being less sensitive to contamination from extracerebral changes in oxy- and deoxy-hemoglobin (Gagnon et al., 2012).

Methods to enhance signal quality based on the anti-correlated relationship between oxy- and de-oxygenated hemoglobin signals (Cui et al., 2010) are widely reported. However, our analysis suggests that applying this technique to fNIRS data provides for no improvement in discriminating between brain states; indeed it reduces performance in three of four of the datasets compared to when oxygenated hemoglobin signals alone are assessed. This indicates that the benefits of applying this algorithm to fNIRS measurements are largely cosmetic and may not provide any objective improvement to signal quality.

By utilizing datasets that measured changes in oxygenation from multiple brain regions, we demonstrated that the oxygenated hemoglobin signal provides the greatest information about neural state related to the stimulus/task compared to de-oxygenated hemoglobin and total hemoglobin. Evaluating the decoding performance from different brain regions is necessary as the systemic signal contribution is not heterogenous over the head, with greater contamination from Mayer waves and pial veins in the motor cortex region (Gagnon et al., 2012). Additionally, the relative contribution of neural activity to the measured signal varies across the head in fNIRS measurements due to variation in skin and skull thickness, and variations in relative depth of the neural tissue of interest. Despite these differences, we observed consistent benefit to using the oxygenated hemoglobin signal alone for decoding between experimental conditions.

## Conclusion

Utilizing multiple datasets, we demonstrate that the fNIRS signal generated by oxygenated hemoglobin provides greater information about the brain state related to the experimental condition than the signal generated by de-oxygenated hemoglobin, regardless of brain region, stimulus, or task. Including manually generated linear and non-linear combinations of the oxy- and de-oxygenated signals did not improve the performance of machine-learning decoding beyond using the oxygenated hemoglobin signal alone. This suggests that the de-oxygenated hemoglobin signal provides no independent information beyond that provided by the oxygenated hemoglobin signal alone. We do not suggest that oxygenated hemoglobin signals only be reported in future; rather, by demonstrating that both signals provide predictive power of the experimental condition above chance level, reporting both signals ideally provides for complimentary findings. However, our data have implications for the interpretation of neuroimaging results, particularly in low signal-to-noise conditions where the two chromophore signals may provide incongruent findings. Additionally, in resource-constrained environments such as brain computer interfaces or neurofeedback applications, it may be advantageous to decode the neural signals based on the oxyhemoglobin signal alone, as it is the most informative fNIRS signal in relation to experimental condition.

## Data and Code availability statement

The code to perform the analysis in this manuscript is available on the MNE-NIRS website (http://mne.tool/mne-nirs) and the data for three of the datasets is publicly available on osf.io and GitHub.

## Ethics statement

All data was collected for this study under the Macquarie University Ethics Application Reference 52020640814625.

